# Spatial extinction date estimation: a novel method for reconstructing spatiotemporal patterns of extinction and identifying potential zones of rediscovery

**DOI:** 10.1101/279679

**Authors:** Colin J. Carlson, Kevin R. Burgio, Tad A. Dallas, Alexander L. Bond

## Abstract

1. The estimation of extinction dates from limited and incomplete sighting records is a key challenge in conservation (when experts are uncertain whether a species has gone extinct) and historical ecology (when the date and mechanism of extinction is controversial).
2. We introduce a spatially-explicit method of interpolating extinction date estimators, allowing users to estimate spatiotemporal surfaces of population persistence from georeferenced sighting data of variable quality.
3. We present the R package spatExtinct, which produces spatially-explicit extinction date surfaces from geolocated sightings, including options for custom randomization schemes to improve accuracy with limited datasets. We use simulations to illustrate the sensitivity of the method to parameterization, and apply the method to identify potential areas where Bachman’s warbler (*Vermivora bachmanii*) might be rediscovered.
4. Our method, and the spatExtinct package, has the potential to help describe and differentiate different drivers of extinction for historical datasets, and could be used to identify possible regions of population persistence for species with an uncertain extinction status, improving on non-spatial or imprecise methods that are currently in use.

## 1 Introduction

Biological extinctions are one of the most fundamental processes in ecology, and despite their significance and pervasiveness, they are often impossible to observe directly. Even if the last known individuals of a species are kept in captivity, uncertainty can still emerge depending on researchers’ confidence that the species is extinct in the wild. Moreover, “sightings” of a species are liable to continue long after a species is presumed extinct, compounding uncertainty and potentially fueling hope of rediscovery. (Carlson *et al.*, 2017b) Every so often, true “Lazarus species” are found after absences of a few centuries (like the Bermuda Petrel, *Pterodroma cahow*) or millions of years (like coelocanths, *Latimeria* spp.) But the majority of species are never rediscovered, presenting conservation biologists with difficult decisions: when should a species be pronounced extinct, and at what point should valuable conservation resources be redirected elsewhere? (Collen *et al.*, 2010; David & Davis, 2017)

A variety of approaches have been developed to address these challenging situations. Some examine the relationship between species’ traits and ecology, and observed extinction rates to estimate the probability of rediscovery (Fisher & Blomberg, 2010; Lee *et al.*, 2017b). Others, which we term extinction date estimators (EDEs), make assumptions about the temporal distribution of sightings leading up to extinction, to estimate the most likely date of extinction and corresponding probability of persistence (Boakes *et al.*, 2015). Over the past two decades, a number of methodological advances have made these methods more powerful and precise (Boakes *et al.*, 2015), and in conservation practice, researchers are encouraged to use these different methods in combination when possible (Akçakaya *et al.*, 2017).

However, one notable limitation of almost all of these methods is that they treat extinction as a single event for an entire species. The total eradication of a species is usually the product of spatially-heterogeneous population declines over time, but spatial tools for reconstructing extinctions are lacking. Spatial kriging is sometimes used to interpolate first and last dates of appearance (Emery-Wetherell *et al.*, 2017), but this basic method has a number of limitations. In particular, modeling based on the last observation is comparatively imprecise, as the extinction date estimation literature shows that the last sighting is usually insufficient to make educated predictions about a species’ true extinction date. Moreover, the kriging approach uses a limited regional subset of data, and makes no inferences based on any of the time series aspects of sighting records. In the context of applying this method to recent (and unconfirmed) extinctions, the kriging approach cannot be used to estimate the probability of persistence, or account for uncertainty in the veracity of sightings.

Species distribution models (SDMs; also called ecological niche models, or ENMs) are another valuable tool for reconstructing the biogeography of extinct species. SDMs conventionally relate occurrence data (sightings) to environmental conditions via some form of regression or machine learning, and make inference about the geographic distribution of species based on their ecological niche. SDMs can be used to reconstruct the shifting distributions over time for extinct species like the megalodon (*Carcharocles megalodon*) in conjunction with extinction date estimators (Pimiento & Clements, 2014; Pimiento *et al.*, 2016), though this application is tenuous over shorter timescales, as most SDM methods assume that distributions are at equilibrium within the scale of modeling. SDMs have also been used with long-extinct species to recover biologically-meaningful information from biogeographic data; for example, a recent study on the Carolina parakeet (*Conuropsis carolinensis*) identified two distinct subspecies’ ranges, and showed that only one subspecies exhibited partial seasonal migration (Burgio *et al.*, 2017). In the shorter term, SDMs are a critical tool for conservation planning, and can be used to help guide the search for possibly-extinct species, even alongside extinction date estimators. (Makenov, 2018) However, for the rarest species, the necessary occurrence data may be impossible to collect. Some workarounds exist, like using data from related species (Dunn *et al.*, 2015), or using Bayesian belief networks to formalize *ad hoc* hypotheses about the species’ niche (Grainger *et al.*, 2017). Even then, environmental suitability may be a poor proxy for presence especially for a species near extinction; in these cases, the total suitable area is likely to be much broader than the true area of occupancy. Methods from the occupancy modeling literature that address this pattern tend to be more data intensive, and require a depth and regularity of observations and abundance data that most putatively- or near-extinct species lack.

We therefore identify a major unaddressed need: researchers interested in reconstructing spatiotemporal patterns of extinction have limited options without explicit data on population declines. While the theoretical underpinnings of extinction date estimators could likely be extended to produce explicitly-spatial analytical approaches, these extensions have yet to be developed, creating an opportunity for the development of a computational, approximate approach. Here, we introduce the idea of *spatial extinction date estimators* (SEDEs) as a tool for recovering the geographic patterns of extinction and identifying where possibly-extant species might be rediscovered. The method we propose uses georeferenced sighting data to estimate extinction dates over a landscape, including for species that have a small chance of persisting somewhere undetected. We show how to optimize the method using simulations, and implement a case study with Bachman’s warbler (Parulidae: *Vermivora bachmanii*), a charismatic North American bird that is likely extinct.

## 2 The Models

### 2.1 Extinction Date Estimators

How do we know if a species is extinct? Extinction date estimators (EDEs) determine the status of a species based on a set of “sightings” including observations, photographs, and physical evidence (like carcasses, specimens, or scat). Sightings can have different levels of support and of validity, and often continue long after a species is extinct. A sighting dataset can be expressed as an ordered set **t** = (*t*_1_, *t*_*n*_), and extinction date estimators make an assumption about the distribution that generates those sightings before (and sometimes after) an extinction event to estimate the true date of extinction *T*_*E*_ (Carlson *et al.*, 2017a). One of the most popular methods, the optimal linear estimator (OLE) is a non-parametric method first used to estimate the extinction date of the dodo (Roberts & Solow, 2003). It assumes that the *k* last few sightings of a species follow a Weibull distribution:

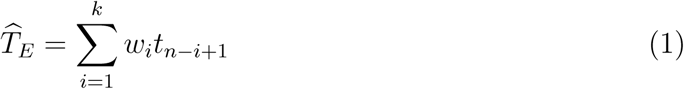

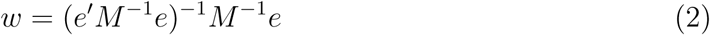

where *e* is a vector of *k* 1’s, and *M* is a *k* by *k* matrix, for which

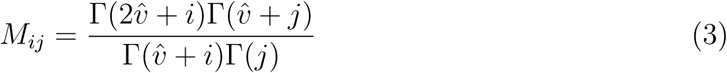

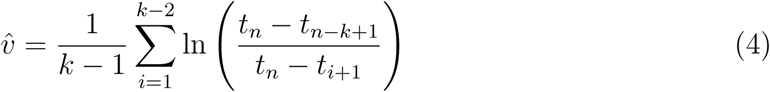

An upper 95% confidence bound is given for the OLE by

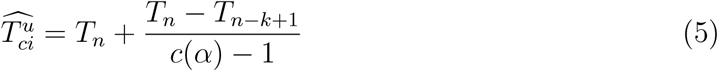

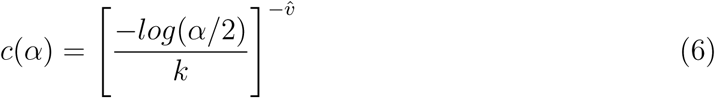

The OLE is one of the most popular EDEs, in large part due to its strong performance even in the face of confounding factors like temporal variation in sampling rates (Rivadeneira *et al.*, 2009). The OLE performs well as an extinction date estimator with limited data, with experimental work finding no universally-optimal sample size and recommending that the method can be best used with all available data (Clements *et al.*, 2013). One notable downside of the OLE method is that it produces extremely wide confidence intervals, especially with larger datasets (Rivadeneira *et al.*, 2009). However, the wide upper confidence bound can be a strength in cases where extreme levels of caution are desired from an extinction date study.

One significant drawback of the OLE, and similar estimators, is that the inclusion of any invalid data proportionally produces significant error in the estimates (Roberts *et al.*, 2010). More recently, a new class of EDEs have been proposed that account for variation in certainty and validity among different sightings (Boakes *et al.*, 2015). These methods tend to be Bayesian, and assume that valid sightings can only exist prior to the extinction date (Solow *et al.*, 2012; Solow & Beet, 2014; Lee *et al.*, 2014). While some of these models can account for variable degrees of confidence in different data sources (Lee *et al.*, 2014), recent work has indicated that expert evaluation of sightings beyond certain and uncertain sightings may be unnecessary (Lee *et al.*, 2017a).

In this study, we adapt two models from Solow & Beet (2014), which assume that data could contain a mix of valid and invalid sightings. The dataset of *n* sightings **t** in an interval (0, *T*] is split in these models into certain (*t*_*c*_, with length *n*_*c*_) and uncertain (*t*_*u*_, with *n*_*u*_ sightings including 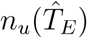 before 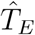) sub-datasets by researchers. The rubric for that split is subjective, but typically, certain sightings involve incontrovertible evidence like a physical specimen, or a clear and uncontroversial photograph. In contrast, uncertain sightings have a much broader range from grainy but plausible video footage (as in the case of the ivory-billed woodpecker), down to unsubstantiated reports of modern-day non-avian dinosaurs. (Smith, 2015) In some cases, it may be helpful for data management to add extra levels of resolution within “uncertain,” such as expert versus novice sightings, even if the model makes no distinction.

If valid sightings occur at a true rate Λ and invalid sightings occur at a true rate Θ, the proportion of valid sightings is given as

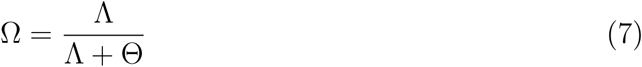

By allowing for a mix of certainty and validity within sightings before extinction, the model makes inferences about the strength of evidence after any hypothesized extinction date. Solow & Beet (SB) model 1 assumes that certain and uncertain sightings follow the same Poisson process. The conditional likelihood of the dataset **t** if the species is extinct is

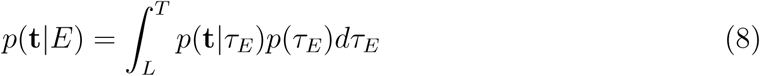

where *t*_*L*_ is the date of the last certain sighting (the starting point of when extinction is possible). Based on the likelihood of the underlying Poisson process for sightings, the likelihood of the dataset given any extinction date is

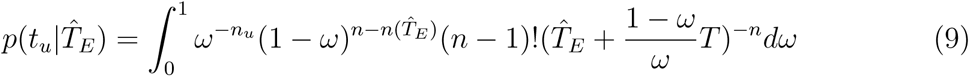

where *ω* is a stand-in for Ω, to allow integration over all possible values of Ω, the true value of which is unknown. In model 2, certain and uncertain sightings are generated by two independent Poisson processes, and the conditional likelihood of the whole dataset is the product of the likelihoods of the respective sub-datasets:

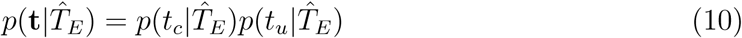

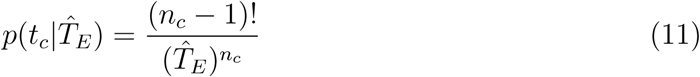

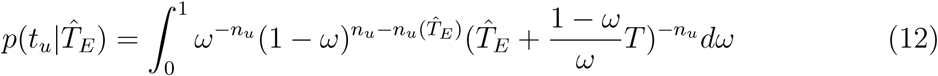

where *ω* is again a stand-in for Ω.

The probability the species went extinct in (0, *T*], an event *E*, can be expressed using Bayes’ theorem:

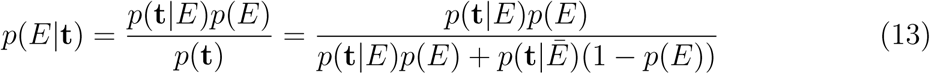

The prior probability of extinction *p*(*E*) is somewhat hard to set, so for explicit calculation, it is often uninformatively set to 0.5 (extinction and persistence are equally likely). If 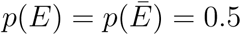, that formula can be reduced to

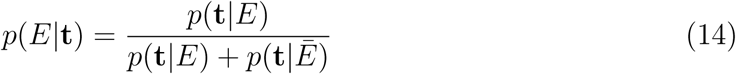

which can be readily interpreted as the “probability of persistence” for the given year. In some cases, researchers may opt instead to use the Bayes factor, which expresses relative support for the alternative hypothesis and is given as

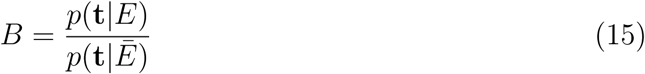

A higher Bayes factor implies stronger support for extinction, where a value of 3 or higher could be taken as strong evidence the species had gone extinct. While the advantage of the Bayes factor is that it avoids the problem of setting *p*(*E*) altogether, both the Bayes factor and *p*(*E|***t**) require setting the conditional likelihood of the data *p*(**t***|E*), which decomposes into

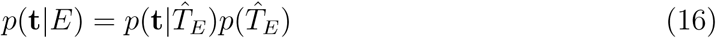

and conversely 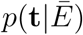 is evaluated using the same function but setting 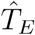 as *T*. The first term can be evaluated as derived above; but the prior probability of a given extinction date 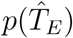 is again subjective and difficult to set. The selection of priors for Bayesian EDEs is an important part of correctly implementing these methods (Solow, 2016); Solow & Beet (2014) suggest either a uniform, linear, or exponential decline after the last certain sighting, and while we have usually elected to use a uniform prior (Carlson *et al.*, 2017b),some researchers may elect to make more informed or constrained choices.

### 2.2 Spatial Extinction Date Estimators

Here we propose a new class of models we term *spatial extinction date estimators* (SEDEs): spatially-explicit interpolations of extinction date estimators using georeferenced sighting data. Whereas EDEs treat extinction as a one-time irreversible event for an entire species, spatial extinction date estimation decomposes extinction into landscape-level extirpation, a set of one-time, irreversible events at the end of local population declines. To do so, our proposed method assigns extinction (extirpation) dates to every cell of a pre-determined grid meant to represent the species’ range or the landscape of interest. In the non-random model, for every grid cell, the *k* nearest neighbor sightings are taken from the centroid, and are run through a specified extinction date estimator. In the random model, a dataset of *N* nearest neighbors are generated for each cell, and estimators are run with a set of *k* records randomly selected without replacement. (Sampling with replacement produces severely distorted estimates, as even one or two extra late sightings can produce centuries-late extinction dates.)

SEDEs use the same sighting datasets as typical EDEs, with the only additional requirement that every record be georeferenced (i.e., sightings are recorded as presence records with date and locality). Depending on data availability, essentially any EDE could be implemented in this modeling framework. Here, we illustrate how SEDEs can be constructed using the optimal linear estimator (OLE), and Solow & Beet’s (2014) Bayesian method for incorporating sighting uncertainty (SB), which we selected based on their ubiquity in the literature, and their demonstrated strong performance relative to other methods. As for non-spatial implementations, the OLE method should only be used for certain sightings (Roberts *et al.*, 2010), while the SB model is designed for use with mixed-certainty data (a common problem in the sighting record of extinct or putatively-extinct species).

In that they reconstruct range contraction over time, SEDEs are, a at least in principle, temporally-dynamic species distribution models: though they use no environmental covariates to make predictions (as “ecological niche models” do), they similarly make a model-based inference about the geographic range of a species based on known occurrence points. For species that are extinct, SEDEs can be used to describe the spatiotemporal process of extinction, which may give clues as to the mechanism. For example, a panzootic disease might spread in a spatial wavefront from a single location (Lips *et al.*, 2008); in contrast, habitat loss or land use change might correlate in a patchwork fashion with local extinctions across a landscape (Preston *et al.*, 2012). In addition to reconstructing the pattern of extinction across a landscape, SEDEs can be used to identify possible zones of persistence for species with an uncertain extinction status. Even if a species-level EDE suggests a low probability of persistence, spatially-subsetted data may indicate potential areas with low support for extinction. This can be done by identifying areas where either the OLE or SB model estimates 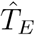 later than the present year. (The confidence intervals for the OLE method could also be used for this purpose, though we discourage this as anything other than an extremely conservative estimate, given how wide these confidence intervals tend to be; see Clements *et al.* [2013]). Additionally, the SB model can also be used to calculate the posterior probability of persistence for a given year, based on a given set of prior assumptions. Zones of potential persistence can then be delineated with a desired confidence level.

## 3 The R Package

We developed the `R` package `spatExtinct` to implement SEDEs for use in historical ecology and conservation research. The package utilizes some pre-existing functions, including the OLE implementation in the package `sExtinct` (Clements, 2013); anand includes some new functions, including an R implementation of Solow & Beet’s model, which has been published (Carlson *et al.*, 2017b) but is streamlined in our package.

### 3.1 Estimating Spatial Extinction Surfaces

The primary function of `spatExtinct` is to spatially interpolate the models we describe above, using spatially-explicit occurrence data. The OLE and SB models are respectively implemented in the `spat.OLE` and `spat.SB` functions. These methods are fairly computationally intensive and work on a cell-by-cell basis, but the package includes options for parallelization and adaptive estimation (with a set convergence threshold to reduce the number of runs). The only required data to run the basic functions are: (1) a data frame with sightings’ decimal longitude and latitude, date (year or any other internally consistent way of denoting time), and scored sighting quality (expert-verified, plausible, or uncertain); and (2) a raster onto which extinction dates can be projected. For the raster, local grids can be used based on the area of interest; or, if researchers are interested in patterns across a species’ entire range, we suggest that species distribution models can be used to define appropriate boundaries for interpolation, as they can be generated from the same sighting data used by the package, and provide an intuitive constraint on outermost area of occupancy. The format in which `spatExtinct` uses data is readily usable by species distribution modeling packages like dismo. (Hijmans *et al.*, 2013)

### 3.2 Estimating Zones of Persistence

The primary utility of `spatExtinct` is estimating the last likely year of presence on a cell-by-cell basis across landscapes. However, there is one readily-obvious extension for species that may *not* be entirely extinct: `spatExtinct` can be used multiple ways to identify potential zones of persistence. This can be done by using spat.OLE and spat.SB to find areas where 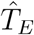 is later than the current year, but we have also included an explicitly probabilistic function spat.SB.probs that estimates the probability of persistence in a set year. That approach would work based on a hypothesis test that the date of extinction *T*_*E*_ is not before the current time *T*, where assuming some significance cutoff *α*, we delineate cells (*i, j*) for which

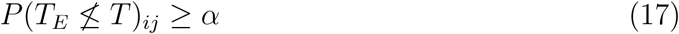

For instance, if we wanted to identify areas where there is at least a 10% chance the species is not yet extinct, we would set *α* = 0.10 and map all cells meeting that criterion. We suggest that this more effective, or at the least more subjective on the user end, than simply cutting off by 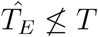.

We also suggest that researchers can easily interface SEDEs and ENMs, as a simple but powerful approach to optimizing rediscovery efforts with almost no *a priori* assumptions. Ecological niche models represent the probability of a species’ presence relative to a given set of environmental variables, and so a combined probability of rediscovery can be conceptualized as

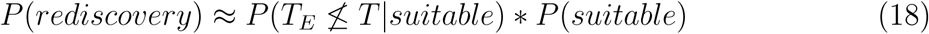

This relative approach does not represent a “true probability” of rediscovery, but can be used to prioritize search efforts in areas of suitable habitat with plausible undiscovered populations, or to guide reserve design for cryptic or rarely-sighted species, for example. The implementation of that process is particularly flexible; for instance, we can identify at least three approaches:

1. Using a thresholded SDM as the base grid for `spat.OLE` or `spat.SB` and identifying zones where 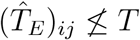
2. Using a thresholded SDM as the base grid for `spat.SB.probs` and identifying zones where *P* (*T*_*E*_ *≰ T*)_*ij*_ *≥ α*
3. Using raw ENM suitability values and `spat.SB.probs` on the same landscape, taking the product to approximate *P* (*rediscovery*), and identifying areas where the combined probability is above either a pre-determined threshold or a quantile (e.g. mapping the top 10% of sites as hotspots of possible rediscovery).

We include no direct tool to interface ENMs and SEDEs, not due to lack of feasibility, but because we stress the importance of careful user-end precision in the implementation of ENMs. All methods for ecological niche modeling are sensitive to sampling bias, pseudoabsence generation, environmental variable set selection, and parameter tuning decisions (Elith & Graham, 2009; Merow *et al.*, 2014); rather than include an automated workflow, we encourage researchers to make careful decisions about these factors when building species distribution models for rare, vulnerable species. We include an example here using these methods to identify possible zones of rediscovery for Bachman’s warbler.

## 4 Applications

We briefly discuss two examples of how our method can be implemented, to illustrate the flexibility of the package, and the relevance in both historical ecology and current conservation work.

### 4.1 Simulations: Validation and Parameterization

We use simulated extinctions to demonstrate the relative accuracy of the methods as a function of data availability and model implementation. Simulations were run on an *n* by *n* square landscape where extinction dates occur on a linear gradient from 5 to 5*n* (e.g. a 10 by 10 landscape has cell extinction dates ranging from 5 to 50). For every year in the interval of [5, 5*n*), the landscape is constrained to areas where populations are extant; *p* “sighting” points are generated on that surface within every year for a total of *p*(5*n-*1) points (see **Figure 1** for an example). As a consequence of this, later extinctions slightly autocorrelate with denser sightings. We used simulated datasets, iterated over adjustable model parameters, to develop an optimization protocol for spatExtinct. We present our analyses with two accuracy measures: correlation of estimated and real extinction surfaces (is the **pattern** clear?), and mean squared error between estimated and real surfaces (how **precise** are extinction date estimates?). More advanced tuning issues are discussed in the Supporting Information, but here, we examine a handful of basic questions:

**Figure 1:**
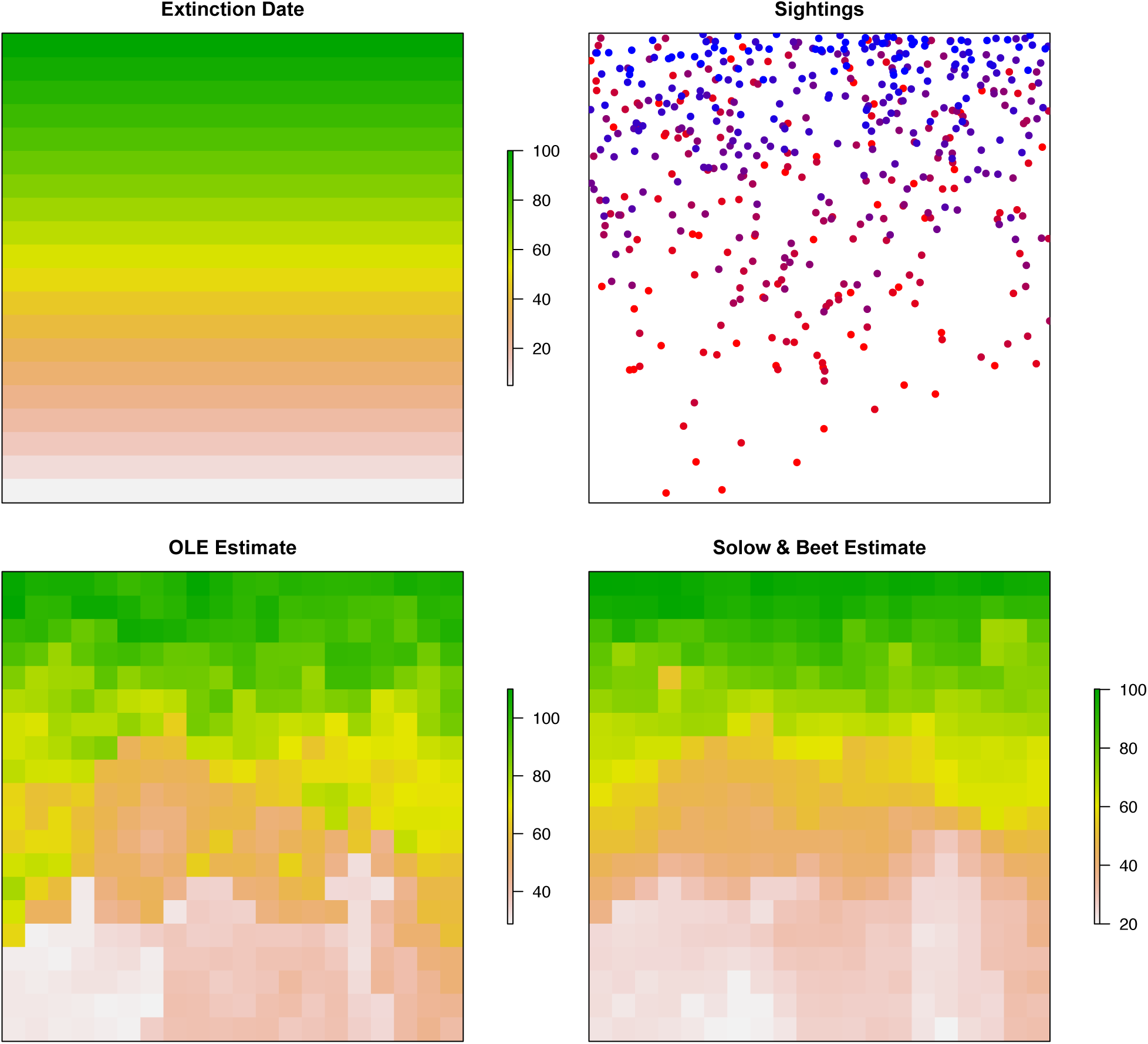
An example simulation and implementation of spatial extinction date estimation. Simulations run with 5 sightings per year over a 20 by 20 landscape. OLE and S&B models parameterized using best practices tuning described in the main text, randomized over 100 iterations.

#### 4.1.1 Which model performs best?

Independent of randomization method, we found that Solow & Beet’s method consistently performed better, with higher correlation coefficients (which were fairly insensitive to parameterization, unlike the OLE method) and lower error rates (**Figure 2**). Consequently, for both pattern estimation and explicit extinction date estimates, the SB method is likely a better one than the OLE (especially if working with mixed-certainty data; see below). However, there are still cases where researchers may want to use the OLE method; most notably, if attempting to delineate regions with a minor chance of persistence, the characteristically wide upper confidence bound of the OLE method (Clements *et al.*, 2013) is a strength of the approach. We suggest the best practice is simply to present both, with necessary disclaimers about known levels of accuracy.

**Figure 2:**
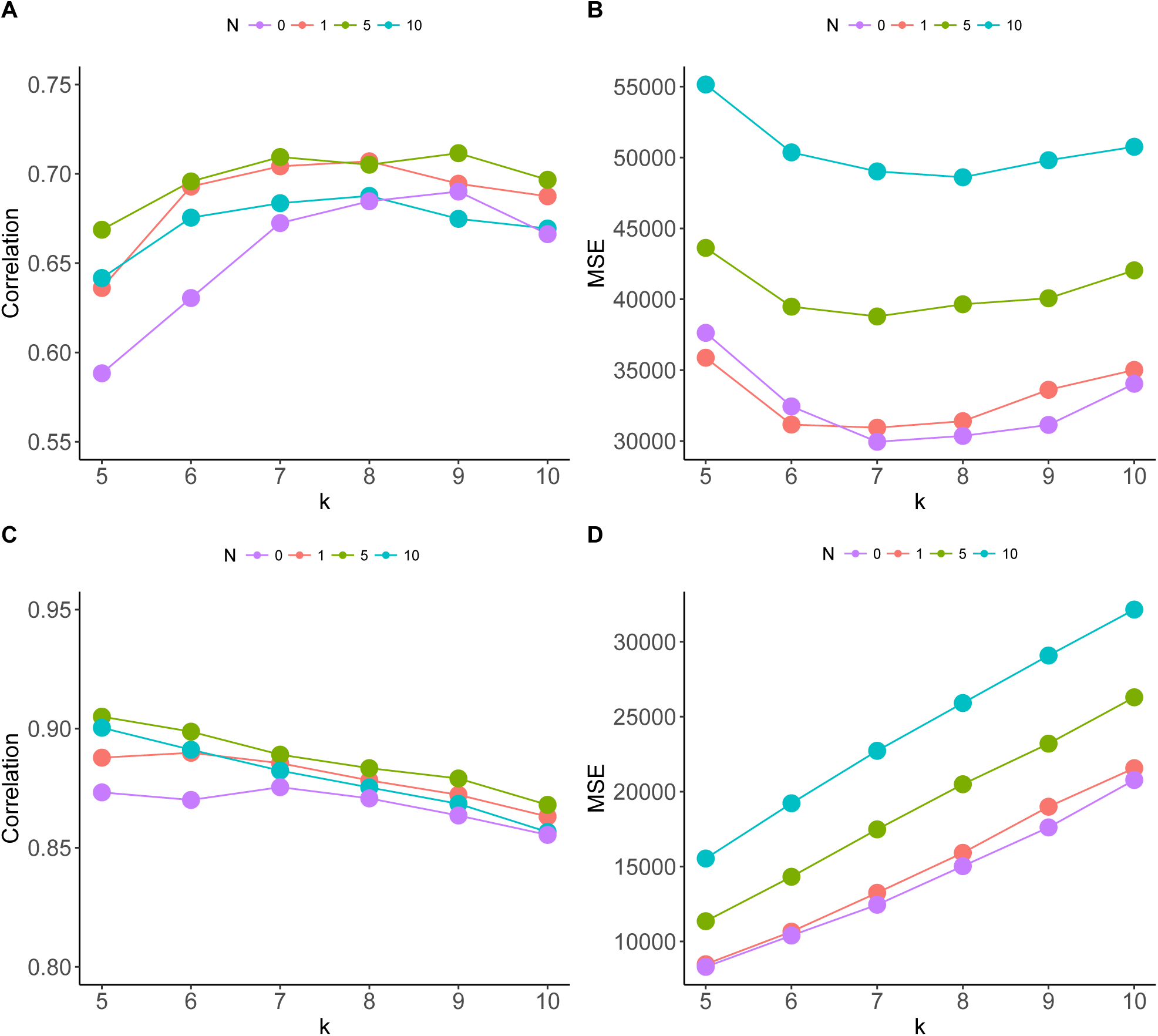
Optimizing over neighborhood size. Accuracy metrics are given for the spatial OLE (A,B) and spatial SB (C,D) models, averaged across 20 simulations for each *k, N* pair. We calculated the average correlation between estimated extinction surfaces and true extinction dates (A,C), and the mean squared error of those estimates (B,D), for different neighborhood sizes (*k*) and levels of randomization (*N*, where “1” indicates *N* = *k* + 1).

#### 4.1.2 How sensitive are models to parameterization?

For the OLE method, we found that error rates were minimized around *k ≈* 7, with limited randomization (*N* = *k* + 1) or no randomization (*N* = *k* + 0); however, an intermediate level of randomization (*N* = *k* + 5) significantly improved correlations with real surfaces (**Figure 2**). For the SB method, smallest levels of *k* minimized error, with increasing levels of randomization similarly leading to decreasing accuracy across *k* values. Again, intermediate levels of randomization (*N* = *k* + 5) maximized correlations, especially at low *k* values. Across methods, we suggest this indicates an unsurprising tradeoff between pattern inference and the precision of estimates: randomization helps smooth estimated surfaces, and makes them easier to interpret, but at the cost of local accuracy. For the most precise estimates (e.g. delineating potential zones of persistence), researchers should select a small neighborhood size and limited or no randomization. In contrast, mild randomization may help researchers accurately interpret the spatial patterns of extinction over landscapes.

#### 4.1.3 How does sample size affect accuracy?

Higher sample sizes consistently improve model performance (**Figure 3**). For both the OLE (A,B) and SB (C,D) method, accuracy plateaus noticeably around 40-50 points. Correlations plateau more substantially, whereas increased sample size continues to reduce estimates’ error. However, we note that the error of SB estimates around 15-25 points is still comparable to that of OLE estimates around 50 or more, highlighting its better performance as an estimator.

**Figure 3:**
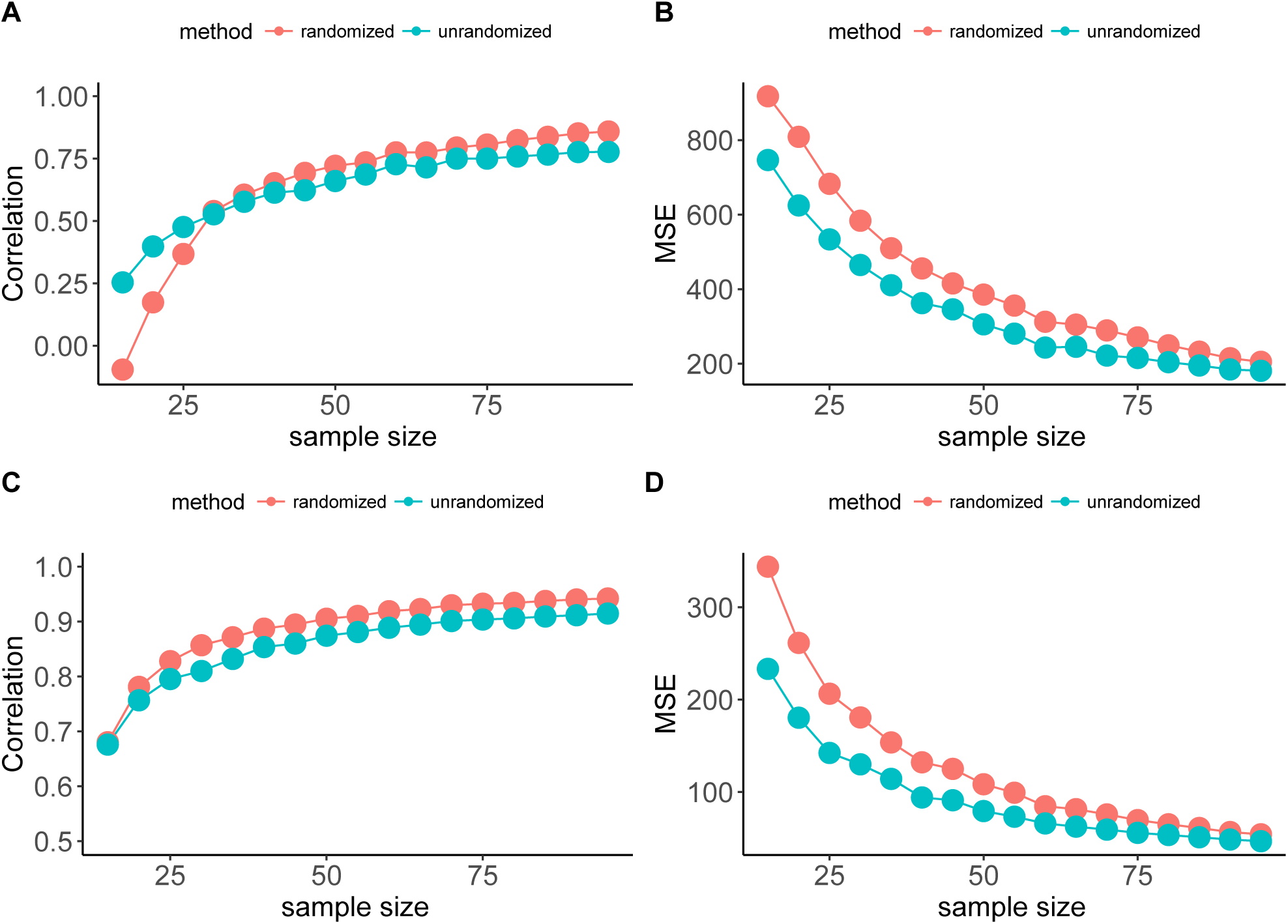
Sample size improves estimates. Results are averaged across 20 simulations for each sample size level (each with 10 resampling instances for samples within a given simulation), with neighborhood size for the OLE (*k* = 7, *N* = 12; A,B) and SB (*k* = 5, *N* = 10; C,D) models taken from the best practices determined in **Figure 2**.

#### 4.1.4 Can mixed certainty sighting data be effectively utilized?

The most significant strength of the Solow & Beet (2014) model is the capacity to use mixed-certainty sighting data, including potentially invalid sightings, to improve extinction date estimates. We show that this benefit still exists when the Solow & Beet models are spatially interpolated (see **Figure 4**); our analyses suggest that a higher number invalid points only severely reduces model performance when valid points have a high uncertainty rate (see **Supporting Information**). However, a quality control option that removes uncertain sightings exists in the OLE functions, for cases where researchers may want to include OLE analyses despite uncertain sightings. This method, while imprecise, still vastly improves the OLE’s performance compared to the dramatic negative impact of inaccurate sightings on unrestricted OLE analyses.

**Figure 4:**
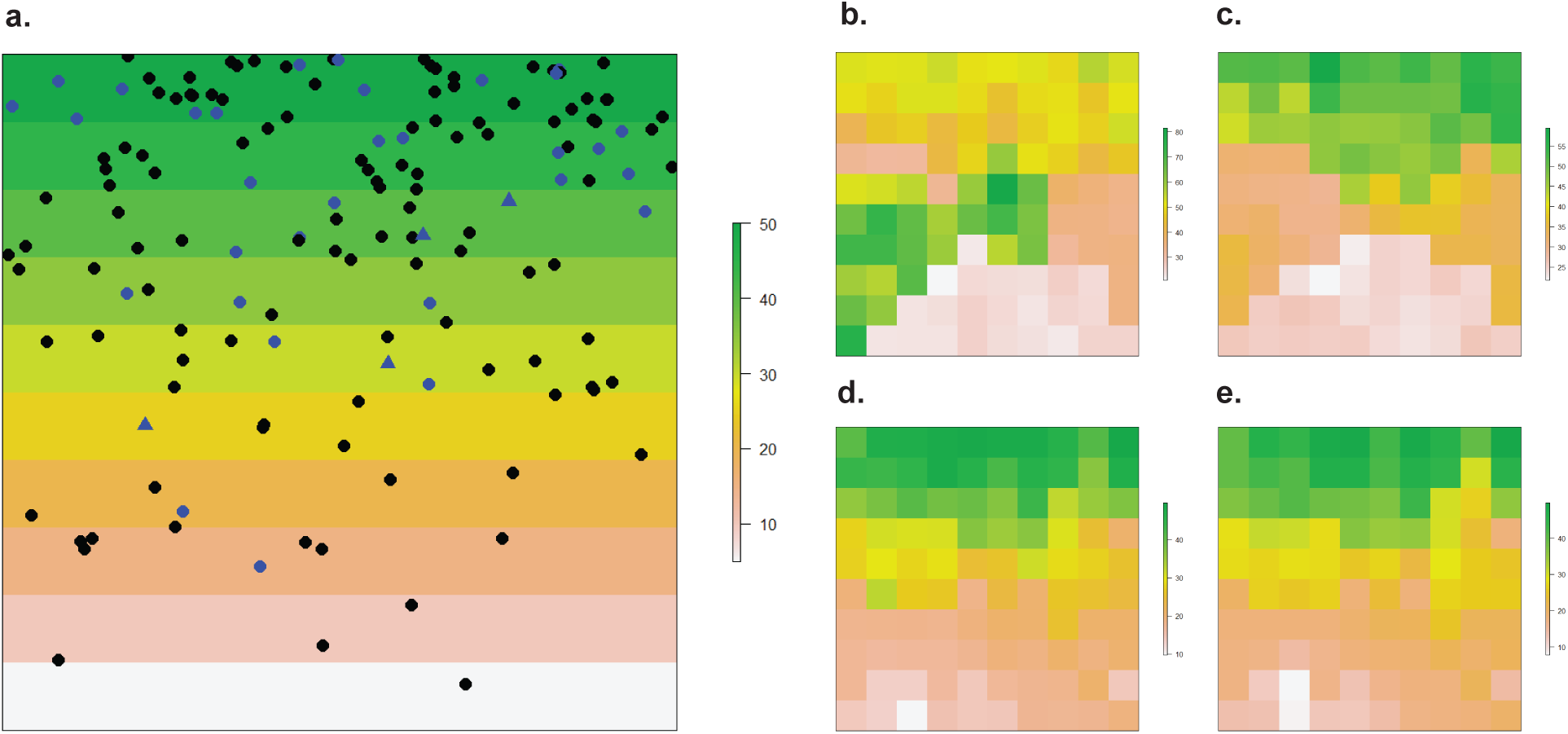
Accounting for mixed-certainty, mixed-accuracy sightings. (A) An example simulated dataset over 50 years; 3 valid sightings are recorded per year (black), 20% of which are recorded as “uncertain” (blue circles), while 4 erroneous sightings are recorded total (blue triangles), with randomly assigned dates. Spatial OLE (*k* = 7, *N* = 12) without quality control (B) is highly prone to misinterpretation, versus spatial OLE with quality control (C), which noticeably improves predictions but also limits data available for use. Solow & Beet’s model (*k* = 5, *N* = 10) includes mixed certainty data and performs noticeably better, regardless of selecting model 1 (D) or model 2 (E).

## 4.2 Case Study: Bachman’s Warbler

To illustrate the potential use of `spatExtinct` for estimating zones of plausible persistence, we provide an example using sightings of Bachman’s warbler, a species that is widely believed extinct despite recent, extremely controversial “sightings.” The species is believed to have experienced sharp population declines between 1910 and 1930, possibly due to habitat loss in the southern United States or its overwintering grounds in Cuba (due in part to hurricane damage); however, as for many species, the precise reasons for decline are controversial and unresolved (Stevenson, 1972; Pimm & Askins, 1995; Huntington & Barbour, 1936; Hamel, 2011).

Unlike higher-profile North American extinct species like the ivory-billed woodpecker (*Campephilus principalis*) or passenger pigeon (*Ectopistes migratorius*), the status of Bachman’s warbler has received comparatively little attention. To the extent of our knowledge, the only modeling study published on the subject suggested an extinction date of 1961 (upper 95% CI: 1967) based on physical evidence only, and 1964 (95% CI: 1967) including expert-opinion sightings (Elphick *et al.*, 2010); however, controversial and unverifiable sightings have been reported as recently as 2001 (Chamberlain, 2003). It is not implausible that Bachman’s warbler could go undetected for several years, given the species’ biology. One naturalist’s report from the early 20th Century notes: “These birds are very hard to detect…in fact I cannot recall a bird that moves as rapidly as Bachman’s Warbler does in the breeding season” (Chamberlain, 2003). On the other hand, recent sightings are particularly dubious given the risk of misidentification, as the species has a strong resemblance to the extant hooded warbler (*Wilsonia citrina*).

To evaluate the status of the species, we assembled a georeferenced dataset containing all readily-accessible known records of Bachman’s warbler in the continental United States. Sightings were sorted into an established set of three categories (Solow & Beet, 2014; Carlson *et al.*, 2017b): confirmed sightings with physical evidence, expert-supported sightings, and unconfirmed but plausible sightings. We developed a species distribution model following a similar protocol to Burgio *et al.* (2017), using the package ENMeval to tune maximum entropy (MaxEnt) species distribution models for the data. We used the data to develop a spatial surface for the probability of persistence in 2017 using the `spat.SB.probs` function in `spatExtinct`, and we use the to estimate the probability of persistence in 2017 based on certain and uncertain sightings. (See **Supporting Information** for more details on data collection and model implementation.)

In total we found 118 usable historical records of Bachman’s warbler (64 certain and verifiable sightings, 46 strongly plausible expert sightings, and 8 implausible or novice sightings). Sightings of Bachman’s warbler dated back as far as the species’ taxonomic description in 1833, and as late as the last certain sighting in 1959, with both sightings recorded near Charleston, South Carolina. The last unconfirmed “sighting” in 2001 was recorded in Congaree Swamp Park in Richland, South Carolina (Chamberlain, 2003). After georeferencing, there were 86 spatiotemporally unique sightings recorded, at 47 unique localities. While this is sufficient for species distribution modeling (Proosdij *et al.*, 2016), it is still astonishing how limited data are for a charismatic North American species that only went (probably) extinct in the last century. (Georeferenced data were limited enough that we suggest interpreting the below analyses as more of a vignette for how a study could be designed using `spatExtinct`, than a definitive appraisal of the status of the species.)

The hypothesis that Bachman’s warbler has been extinct for several years was supported by both the optimal linear estimator (all data: 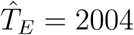, 95% CI = (2001,2012);certain sightings only: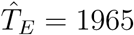, 95% CI = (1959,1987)) and the Solow & Beet method (model 1: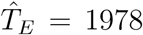, *p*(*T*_*E*_ *≥* 2017) = 0.011; model 2: 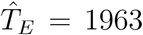, *p*(*T*_*E*_ *≥* 2017) =1.5 *×* 10^−5^). Without any spatial information, these results suggest the species can be safely presumed extinct in 2018; in fact, these estimates suggest the species was probably already extinct by 1967, when it was included on the first federal listing of endangered species in the United States (the “Class of ’67”). The species is still listed despite its apparent extinction; in 2015, the U.S. Fish and Wildlife Service most recently reassessed the species, maintaining its Endangered status and explaining, “We considered recommending delisting Bachmans warbler; however, because Bachmans warbler is difficult to detect and identify (Chamberlain 2003) and the lack of formal extensive search efforts over the last 27 years, considerable uncertainty remains as to its status.” (Sisson, 2015)

Our species distribution model agreed with prior expert knowledge, suggesting a broad geographic division between a coastal Atlantic range, and an inland range following the Mississippi and Ohio River basins, tracking the distribution of baldcypress (*Taxodium distichum*) in the South. Previous work suggested these ranges were continuous and connected above the Gulf of Mexico, though our model suggests the ranges may have been more separate (Hamel, 2011). The spatial Solow & Beet model suggested that, despite one late uncertain sighting in northern Louisiana, the western range of the species was likely gone by the early 20^*th*^ century (**Figure 5**). The model suggested persistence in the 1960s or 1970s along the coast of Georgia and South Carolina, as well as (surprisingly) around the Chesapeake Bay. These estimates still suggest that the species was likely extinct by the mid-1970s.

**Figure 5:**
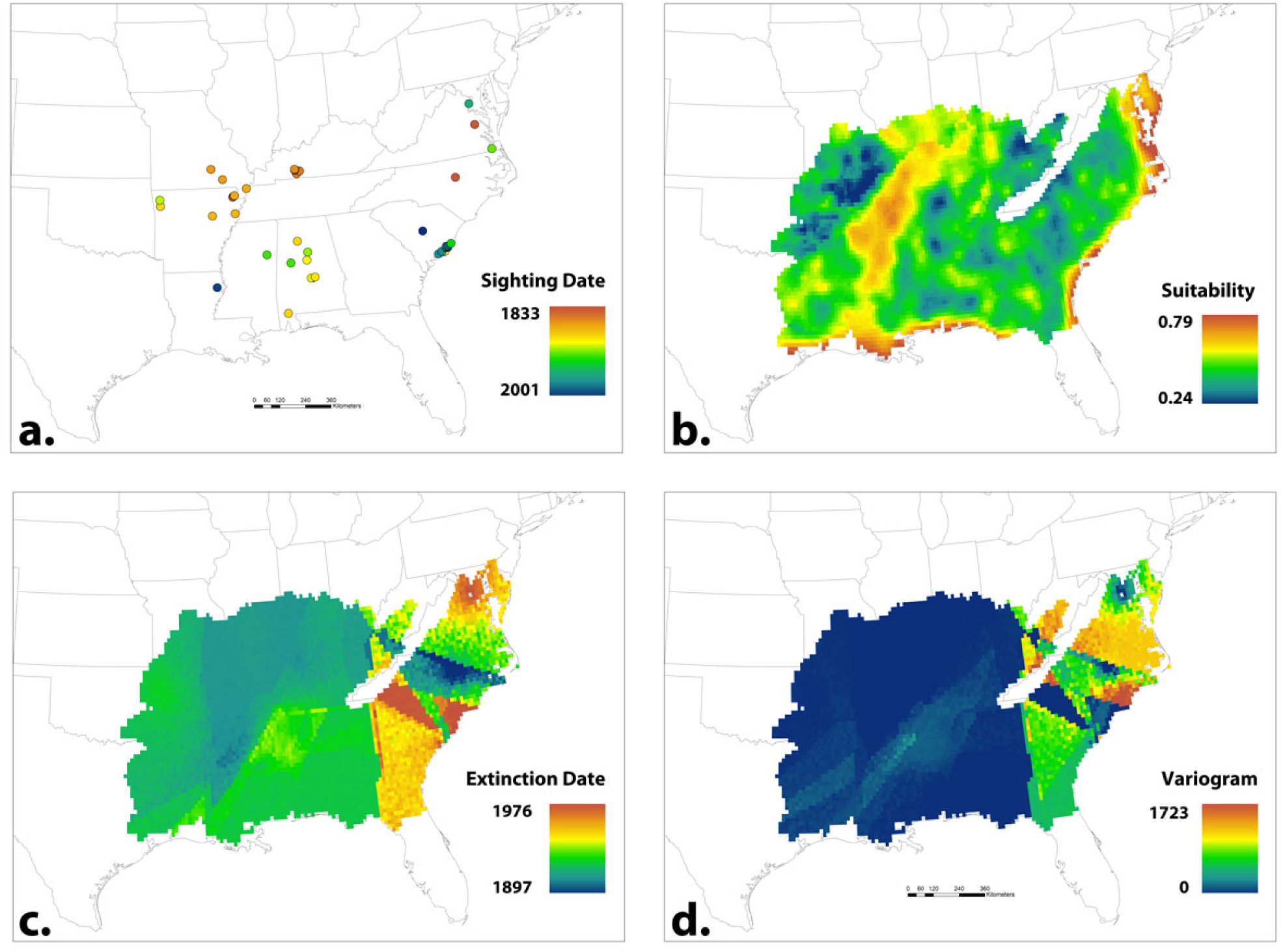
Mapping Bachman’s Warbler with limited occurrence data. (A) Sighting records (*n* = 118). (B) MaxEnt ecological niche model. (C) Extinction date reconstructed using Solow & Beet’s model 2, on recommended best practices settings. (D) Variance in Solow & Beet estimates (higher variance means more uncertain estimates).

As a final test, we used the probability of persistence (*p*(*T*_*E*_ *≰* 2017), generated with the `spat.SB.probs` function) to test the hypothesis of local extinction and verify the minimal chance of rediscovery. In some of the most uncertain and probably error-prone areas (West Virginia and North Carolina) the probability was surprisingly high. However, the model still found broad regions with a small but nontrivial (5-20%) posterior probability of presence. Using the species distribution model as an additional probability filter, we found three major areas of potential rediscovery: coastal Georgia and South Carolina, the Mississippi delta in Louisiana, and a broad patch from North Carolina up through Maryland (**Figure 6**).

**Figure 6:**
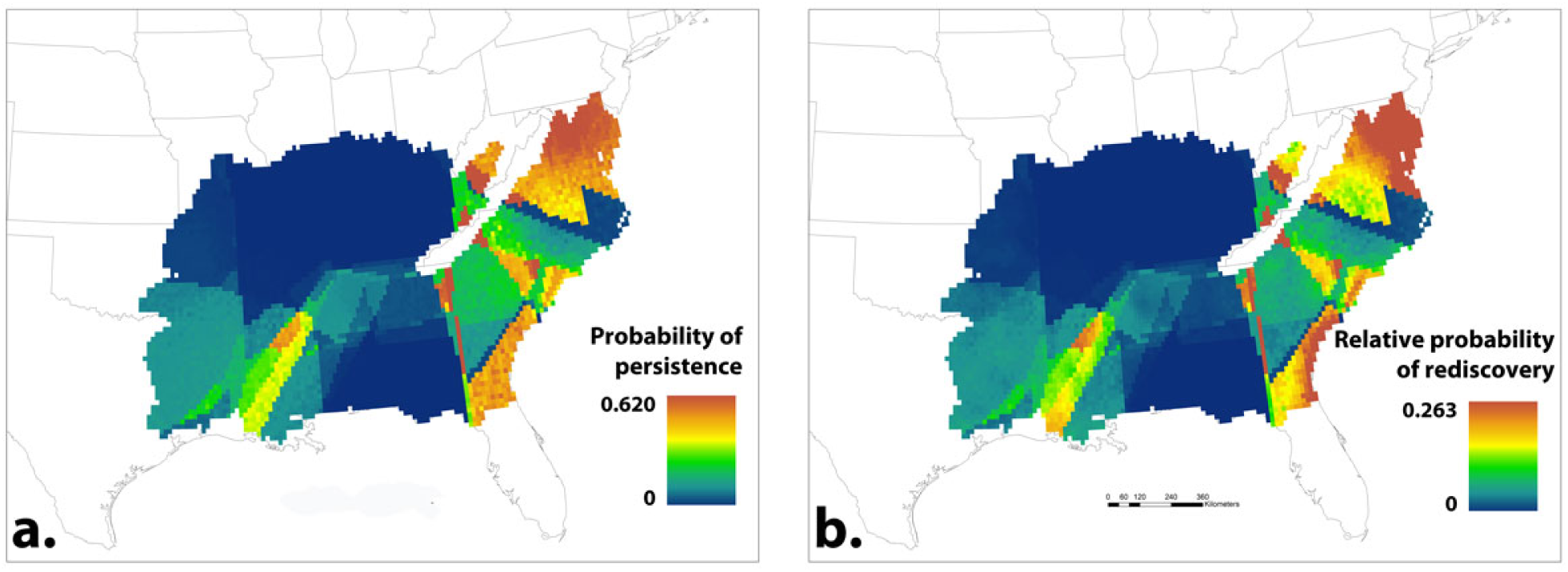
Where are the odds of rediscovery highest? (A) Probability of persistence based on Bayesian hypothesis testing in Solow & Beet model (extinction date only). (B) Combined probability of rediscovery (product of probability of persistence from EDE and environmental suitability from ENM).

The idea that there might be undiscovered, isolated populations of Bachman’s warbler in the southeastern United States—especially in South Carolina—is far from new. Official U.S. Fish & Wildlife Service documentation still lists the species as having a range along coastal South Carolina and the southernmost end of Florida (the latter being part of its migratory range; USFWS [2018]). In the mid-1970s, a series of unconfirmed sightings were reported in the I’on Swamp in coastal South Carolina. (Shuler, 1976) After unconfirmed but plausible sightings in 2000 and 2001 in Congaree National Park, the park was surveyed in 2002 without any success (Chamberlain, 2003), suggesting that even if there were late persistent populations in the area, they are likely gone by today.

The prediction of a possible zone of persistence further north is surprising, given the absence of late plausible sightings, but could possibly merit further investigation. Most promising, in our opinion, is the possible zone of persistence in the Mississippi basin. While the swamps of Congaree have been searched, remaining baldcypress swamps along the Mississippi could just as plausibly harbor an undiscovered remnant population; the 2015 FWS assessment noted optimistically that efforts to rediscover the ivory-billed wood-pecker in similar habitats might incidentally lead to the warbler’s rediscovery. (Sisson, 2015) Given that Bachman’s warbler was inconspicuous at best, it may also be possible that the bird, if it still persists, will be found in unexpected areas. SDMs have been proven useful in finding new populations of other elusive species (Menon *et al.*, 2010; Williams *et al.*, 2009; Fois *et al.*, 2015), so it may well be worth looking in the areas suggested by our models. But all evidence suggests the species is likely extinct, and the decision to keep searching for Bachman’s warbler may also detract from other, more promising conservation efforts—and the decision to continue the search depends at least as much on values as it does on model results. (Akçakaya *et al.*, 2017; Carlson *et al.*, 2017b; David & Davis, 2017; Jackson, 2006)

## 5 Discussion

Here, we have introduced the idea of a spatially-explicit approach to extinction date estimation, and an associated R package. Spatial extinction date estimation makes an important conceptual link between extinction date estimation and species distribution modeling, and when parameterized correctly, we suggest that our method can be readily used to reconstruct the spatial pattern of range loss during a species’ extinction. Provided that sufficient data is available, we suggest that spatial extinction date estimation has tremendous promise as a new tool for assessing the status of putatively-extinct species like Bachman’s warbler, even when some sightings are suspected to be invalid.

Spatial extinction date estimation is a new method, and our simulations suggest it works fairly well. However, the computational approach to approximation we use in the `spatExtinct` package would likely be outperformed by an analytic approach that adapts a model like Solow & Beet’s to explicitly consider spatial autocorrelation in extinction dates and spatial heterogeneity in sighting rates. (In fact, we hypothesize that much of the temporal heterogeneity in sighting rates that these models account find could be attributed to the combination of spatial heterogeneity and species’ contracting ranges.) We encourage the development of superior modeling approaches, and similarly encourage caution interpreting the results of any model. Users of the method, and the R package, should carefully consider the bias different models contribute: for example, Solow & Beet’s model 1 is strongly influenced by the last uncertain sighting and predicts persistence more commonly than model 2, which is more strongly influenced by the last certain sighting (Kodikara *et al.*, 2018). The relative accuracy of the two models is influenced by the proportion of uncertain sightings that are invalid, something users of our method should strongly consider.

After the extinction of a species, the amount of biological information that can be recovered rapidly declines, especially for poorly-documented recent extinctions without a detailed specimen or fossil record. As the sixth mass extinction accelerates due to forces like climate change, an increasing amount of information on global biodiversity is irretrievably lost (Barnosky *et al.*, 2011; Ceballos *et al.*, 2015; Urban, 2015). In particularly data-deficient situations, the mechanism of an extinction may be uncertain. In some cases, a species’ status as extinct may itself be uncertain. Spatially-interpolated extinction date estimators, and their implementation in the `spatExtinct` package, are designed to address both of these data-limited situations. We suggest, though, that the strongest use of SEDEs is alongside other tools. Statistical methods like ENMs hold clear promise as a companion to SEDEs, but sighting data can only accomplish so much. Using these tools alongside specimen work, however, is likely to be especially powerful as a path of inquiry. Identifying cause of death from specimens may help explain mortality patterns and develop informative population models (Cunningham & Daszak, 1998), and stable isotope work can help recover key information about changing patterns of diet or migration (Hilderbrand *et al.*, 1996). All of these methods in conjunction can help develop a more robust perspective on extinction as a spatiotemporal process, rather than a single event in time, benefiting work in historical ecology and conservation biology alike.

## Acknowledgements

We thank Andrew Beet for the original Matlab code used in Solow & Beet (2014); Adam Butler (Biomathematics and Statistics Scotland) for translating the Matlab code into R; Christopher Clements for helpful feedback and brainstorming; Nathaniel J. Ruess, for moral support the last six years; and Robert Rankin for statistics advice.

## Author Contributions

CJC, KRB, and ALB conceived of the idea for the study. CJC and KRB developed the modeling framework, and CJC developed the R package. TAD and CJC developed the simulations. KRB collected the Bachman’s warbler dataset, and CJC and KRB ran subsequent analyses. All authors contributed to the writing and editing of the manuscript.

## Data Accessibility

spatExtinct can be downloaded from https://github.com/cjcarlson/spatExtinct.Code to reproduce simulation analyses, and the Bachman’s warbler dataset, are available on Figshare at (**link forthcoming**).

## References

Akçakaya, H., Keith, D.A., Burgman, M., Butchart, S.H., Hoffmann, M., Regan, H.M., Harrison, I. & Boakes, E. (2017) Inferring extinctions III: A cost-benefit framework for listing extinct species. Biological Conservation, p. in press.

Barnosky, A.D., Matzke, N., Tomiya, S., Wogan, G.O., Swartz, B., Quental, T.B., Marshall, C., McGuire, J.L., Lindsey, E.L., Maguire, K.C. et al. (2011) Has the Earth’s sixth mass extinction already arrived? Nature, 471, 51–57.

Boakes, E.H., Rout, T.M. & Collen, B. (2015) Inferring species extinction: the use of sighting records. Methods in Ecology and Evolution, 6, 678–687.

Burgio, K.R., Carlson, C.J. & Tingley, M.W. (2017) Lazarus ecology: Recovering the distribution and migratory patterns of the extinct Carolina parakeet. Ecology and Evolution, 7, 5467–5475.

Carlson, C., Burgio, K., Dallas, T. & Getz, W. (2017a) The mathematics of extinction across scales: from populations to the biosphere. PeerJ PrePrints.

Carlson, C.J., Bond, A.L. & Burgio, K.R. (2017b) Estimating the extinction date of the thylacine with mixed certainty data. Conservation Biology, p. in press.

Ceballos, G., Ehrlich, P.R., Barnosky, A.D., García, A., Pringle, R.M. & Palmer, T.M. (2015) Accelerated modern human–induced species losses: Entering the sixth mass extinction. Science Advances, 1, e1400253.

Chamberlain, D. (2003) Additional notes on Bachman’s Warbler. Chat, 67, 5–l0.

Clements, C. (2013) sExtinct: Calculates the historic date of extinction given a series of sighting events. R package version 1.1.

Clements, C.F., Worsfold, N.T., Warren, P.H., Collen, B., Clark, N., Blackburn, T.M. & Petchey, O.L. (2013) Experimentally testing the accuracy of an extinction estimator: Solow’s optimal linear estimation model. Journal of Animal Ecology, 82, 345–354.

Collen, B., Purvis, A. & Mace, G.M. (2010) Biodiversity research: When is a species really extinct? Testing extinction inference from a sighting record to inform conservation assessment. Diversity and Distributions, 16, 755–764.

Cunningham, A.A. & Daszak, P. (1998) Extinction of a species of land snail due to infection with a microsporidian parasite. Conservation Biology, 12, 1139–1141.

David, W.M. & Davis, R.A. (2017) Hopeful monsters—in defence of quests to rediscover long-lost species. Conservation Letters, 10, 382–383.

Dunn, J.C., Buchanan, G.M., Cuthbert, R.J., Whittingham, M.J. & Mcgowan, P.J. (2015) Mapping the potential distribution of the critically endangered himalayan quail *Ophrysia superciliosa* using proxy species and species distribution modelling. Bird Conservation International, 25, 466–478.

Elith, J. & Graham, C.H. (2009) Do they? How do they? WHY do they differ? On finding reasons for differing performances of species distribution models. Ecography, 32, 66–77.

Elphick, C.S., Roberts, D.L. & Reed, J.M. (2010) Estimated dates of recent extinctions for north american and hawaiian birds. Biological Conservation, 143, 617–624.

Emery-Wetherell, M.M., McHorse, B.K. & Davis, E.B. (2017) Spatially explicit analysis sheds new light on the pleistocene megafaunal extinction in north america. Paleobiology, pp. 1–14.

Fisher, D.O. & Blomberg, S.P. (2010) Correlates of rediscovery and the detectability of extinction in mammals. Proceedings of the Royal Society of London B: Biological Sciences, p. rspb20101579.

Fois, M., Fenu, G., Lombrana, A.C., Cogoni, D. & Bacchetta, G. (2015) A practical method to speed up the discovery of unknown populations using species distribution models. Journal for Nature Conservation, 24, 42–48.

Grainger, M.J., Ngoprasert, D., Mc Gowan, P.J. & Savini, T. (2017) Informing decisions on an extremely data poor species facing imminent extinction. Oryx, pp. 1–7.

Hamel, P.B. (2011) Bachman’s Warbler (Vermivora bachmanii), version 2.0. Cornell Lab of Ornithology, Ithaca, NY.

Hijmans, R., Phillips, S., Leathwick, J. & Elith, J. (2013) dismo. Species distribution modeling. R package version 0. 7-23. 2012.

Hilderbrand, G.V., Farley, S.D., Robbins, C.T., Hanley, T.A., Titus, K. & Servheen, C. (1996) Use of stable isotopes to determine diets of living and extinct bears. Canadian Journal of Zoology, 74, 2080–2088.

Huntington, J. & Barbour, T. (1936) The birds at Soledad, Cuba, after a hurricane. Auk, 53, 436–437.

Jackson, J.A. (2006) Ivory-billed Woodpecker (*Campephilus principalis*): Hope, and the interfaces of science, conservation, and politics. Auk, 123, 1–15.

Kodikara, S., Demirhan, H. & Stone, L. (2018) Inferring about the extinction of a species using certain and uncertain sightings. Journal of theoretical biology, 442, 98–109.

Lee, T.E., Bowman, C. & Roberts, D.L. (2017a) Are extinction opinions extinct? PeerJ, 5, e3663.

Lee, T.E., Fisher, D.O., Blomberg, S.P. & Wintle, B.A. (2017b) Extinct or still out there? disentangling influences on extinction and rediscovery helps to clarify the fate of species on the edge. Global change biology, 23, 621–634.

Lee, T.E., McCarthy, M.A., Wintle, B.A., Bode, M., Roberts, D.L. & Burgman, M.A. (2014) Inferring extinctions from sighting records of variable reliability. Journal of Applied Ecology, 51, 251–258.

Lips, K.R., Diffendorfer, J., Mendelson III, J.R. & Sears, M.W. (2008) Riding the wave: reconciling the roles of disease and climate change in amphibian declines. PLoS biology, 6, e72.

Makenov, M. (2018) Extinct or extant? A review of dhole (*Cuon alpinus* Pallas, 1811) distribution in the former USSR and modern Russia. Mammal Research, pp. 1–9.

Menon, S., Choudhury, B.I., Khan, M.L. & Peterson, A.T. (2010) Ecological niche modeling and local knowledge predict new populations of *Gymnocladus assamicus* a critically endangered tree species. Endangered Species Research, 11, 175–181.

Merow, C., Smith, M.J., Edwards, T.C., Guisan, A., McMahon, S.M., Normand, S., Thuiller, W., Wüest, R.O., Zimmermann, N.E. & Elith, J. (2014) What do we gain from simplicity versus complexity in species distribution models? Ecography, 37, 1267–1281.

Pimiento, C. & Clements, C.F. (2014) When did *Carcharocles megalodon* become extinct? a new analysis of the fossil record. PLoS One, 9, e111086.

Pimiento, C., MacFadden, B.J., Clements, C.F., Varela, S., Jaramillo, C., Velez-Juarbe, J. & Silliman, B.R. (2016) Geographical distribution patterns of *Carcharocles megalodon* over time reveal clues about extinction mechanisms. Journal of Biogeography, 43, 1645–1655.

Pimm, S.L. & Askins, R.A. (1995) Forest losses predict bird extinctions in eastern north america. Proceedings of the National Academy of Sciences, 92, 9343–9347.

Preston, K.L., Redak, R.A., Allen, M.F. & Rotenberry, J.T. (2012) Changing distribution patterns of an endangered butterfly: Linking local extinction patterns and variable habitat relationships. Biological Conservation, 152, 280–290.

Proosdij, A.S., Sosef, M.S., Wieringa, J.J. & Raes, N. (2016) Minimum required number of specimen records to develop accurate species distribution models. Ecography, 39, 542–552.

Rivadeneira, M.M., Hunt, G. & Roy, K. (2009) The use of sighting records to infer species extinctions: an evaluation of different methods. Ecology, 90, 1291–1300.

Roberts, D.L., Elphick, C.S. & Reed, J.M. (2010) Identifying anomalous reports of putatively extinct species and why it matters. Conservation Biology, 24, 189–196.

Roberts, D.L. & Solow, A.R. (2003) Flightless birds: when did the dodo become extinct? Nature, 426, 245–245.

Shuler, J. (1976) Three recent sight records of Bachman’s warbler. Chat, 41, 11–12.

Sisson, P. (2015) Bachman’s Warbler (*Vermivora bachmanii*) 5-Year Review: Summary and Evaluation. Technical report, U.S. Fish and Wildlife Service.

Smith, J.F. (2015) The Inclusion of False, Falsified, and Falsifiable Data that Favor an Evolutionary Worldview in the High School Science Curriculum of Public and Private Schools in the Philippines. Christian Perspectives in Education, 8, 2.

Solow, A., Smith, W., Burgman, M., Rout, T., Wintle, B. & Roberts, D. (2012) Uncertain sightings and the extinction of the ivory-billed woodpecker. Conservation Biology, 26, 180–184.

Solow, A.R. (2016) On the prior distribution of extinction time. Biology Letters, 12, 20160089.

Solow, A.R. & Beet, A.R. (2014) On uncertain sightings and inference about extinction. Conservation Biology, 28, 1119–1123.

Stevenson, H.M. (1972) The recent history of Bachman’s Warbler. The Wilson Bulletin, 84, 344–347.

Urban, M.C. (2015) Accelerating extinction risk from climate change. Science, 348, 571–573.

USFWS (2018) Species Profile for Bachman’s warbler (Vermivora bachmanii).

Williams, J.N., Seo, C., Thorne, J., Nelson, J.K., Erwin, S., OBrien, J.M. & Schwartz, M.W. (2009) Using species distribution models to predict new occurrences for rare plants. Diversity and Distributions, 15, 565–576.

